# Leukocyte immunoglobulin-like receptor A5 functions to induce ROS production on innate immune cells

**DOI:** 10.1101/2025.03.16.643524

**Authors:** Zuyi Fu, Matevž Rumpret, Irina Kube-Golovin, Mykola Lyndin, Vera Solntceva, Yuxi Zhao, Anastasia Konieva, Na Liu, Adrian T. Press, Stefanie B. Flohé, Michael Bauer, Gunther Wennemuth, Bernhard B. Singer, Alex J. McCarthy

## Abstract

Activating immune receptors provide mechanisms for phagocytes to elicit important effector functions that promote the killing of microbes. Leukocyte immunoglobulin-like receptor A5 (LILRA5) an orphan immune receptor expressed by human phagocytes and co-localising with FcRγ, remains poorly characterised. To address this, we developed a highly-specific anti-LILRA5 monoclonal antibody that has agonistic properties. Using a specific anti-LILRA5 antibody, we show LILRA5 expression on naïve monocytes and neutrophils, and that ligation of LILRA5 stimulates ROS production. While increased *LILRA5* transcript copy numbers have been associated with sepsis, we also observed increased levels in patients with systemic infection but without sepsis complications. *Ex vivo* bacterial infection of whole blood did not alter surface LILRA5 expression, but LPS stimulation changed expression levels. This indicates that surface LILRA5 expression is dynamic and likely regulated posttranscriptionally, changing responses to different stimuli or over time. Soluble (s)LILRA5 was enhanced in sera from sepsis patients and in supernatants of monocytes that were LPS-stimulated, indicative that shedding of LILRA5 from cell surfaces or that expression of sLILRA5 isoforms provides a mechanism to regulate surface LILRA5 expression levels. Finally, we show that altered surface LILRA5 expression influences LILRA5-induced ROS production capacity. We propose that LILRA5 is a dynamically regulated activating receptor expressed on phagocytes that can stimulate ROS production.

## Introduction

Phagocytic immune cells such as neutrophils, monocytes and macrophages play a key role in the innate immune response to infection (1). They can kill invading pathogens by engulfing them through phagocytosis, releasing antimicrobial molecules and generating reactive oxygen species (ROS). The capacity of these immune cells to respond to stimuli in the environment is attributed to the expression of immune receptors on the cell surface that detect signs of infection. Given the importance of phagocytosis, degranulation and ROS generation, these processes can be activated via several innate immune receptors. This includes receptors containing immunoreceptor tyrosine-based activation motif (ITAM) in the cytoplasmic tail (2–4), or those that co-associate with the ITAM-containing FcRγ (1,5). Though several innate immune receptors co-associate with FcRγ (6), their capacity to activate phagocytes, induce degranulation and/or stimulate ROS production is less well understood.

Leukocyte immunoglobulin-like receptor A5 (LILRA5, ILT11, LIR9, CD85f) is a type I transmembrane receptor and member of the leukocyte immunoglobulin-like receptor (LILR) family (7). *LILRA5* transcripts are expressed by neutrophils, monocytes and macrophages (8,9). Though surface LILRA5 expression has been reported for monocytes and macrophages (8), the expression of LILRA5 on neutrophil surfaces remains unclear (10). Structurally, LILRA5 contains an extracellular region formed of two immunoglobulin (Ig) domains, a transmembrane region that contains a positively charged arginine residue for co-localisation with the ITAM-containing FcRγ and a short cytoplasmic tail (8). It can also be expressed in a soluble form known as sLILRA5 (8,9). Though the ligands of LILRA5 are unknown (7,11), cross-linking LILRA5 on monocyte surfaces induces association with FcRγ, phosphorylation of the ITAMs in FcRγ, increased activation of the Src and Syk kinases and release of pro-inflammatory cytokines (8). Mice with deletion of the *LILRA5* gene displayed an increased susceptibility to bacterial keratitis and a dysregulated inflammatory response (12).

Due to the scarcity of highly-specific anti-LILRA5 mAbs and no knowledge on endogenous and exogenous LILRA5 ligands (10), the expression and functions of LILRA5 are poorly elucidated. Recent reports have shown that *LILRA5* transcripts are significantly increased in human bacterial keratitis and in human sepsis (12,13). This is suggestive that LILRA5 has a role in bacterial defence and could be a useful biomarker for rapid diagnosis of inflammation triggered by bacterial pathogens. In this study, we developed a highly-specific antibody against LILRA5 as a tool to investigate expression and functions. We investigated *LILRA5* transcript levels and surface LILRA5 expression on phagocytes in health and disease, and assessed whether cross-linking LILRA5 could trigger ROS production and phagocytosis.

## Results

### Development of LILRA5-specific mAb with agonistic properties

To analyse the expression of LILRA5 on human immune cells, we developed a mouse monoclonal antibody that specifically recognised LILRA5. To do this, recombinant (r)LILRA5 was expressed in a eukaryotic expression system and purified (**Fig. 1A**). Anti-LILRA5 P4-11A mAb was purified from a hybridoma derived from fusion of myeloma NS1/0 cells with spleen cells from Balb/c mice immunised with rLILRA5-His. We characterised the specificity of the mAb for LILRA5 by testing its binding to Dynabeads (DB) coated with rLILR-His or rLAIR1-His (**Supp. Fig. 1A**). Flow cytometry analysis revealed that anti-LILRA5 P4-11A mAb binds to human rLILRA5, but to not any of the closely related human rLILR that are most likely to be cross-reactive, nor to an immune receptor from a different family called LAIR1 (**Fig. 1B, Supp. Fig. 1B**). The isotype matched IgG did not bind to rLILRA5. Furthemore, anti-LILRA5 P4-11A mAb is bound to rLILRA5-coated DB in a concentration-dependent manner (**Supp. Fig. 1C**). This data suggests that anti-LILRA5 P4-11A mAb has high-specificity for detecting LILRA5. We next investigated the capacity of anti-LILRA5 P4-11A mAb to bind LILRA5 presented at the cell surface. Firstly, we generated a LILRA5+ U937 cell line by transducing the parental U937 cell line with a lentiviral vector encoding the coding domain sequence of *LILRA5* and selecting transductants. We then tested the binding of anti-LILRA5 P4-11A mAb to wildtype U937 and LILRA5+ U937 cell lines. Flow cytometry analysis revealed enhanced binding of anti-LILRA5 P4-11A mAb to the LILRA5+ U937 cells compared to control U937 cells (**Fig. 1C**), revealing anti-LILRA5 mAb P4-11A can specifically detect cell surface expressed LILRA5.

**Figure 1:**
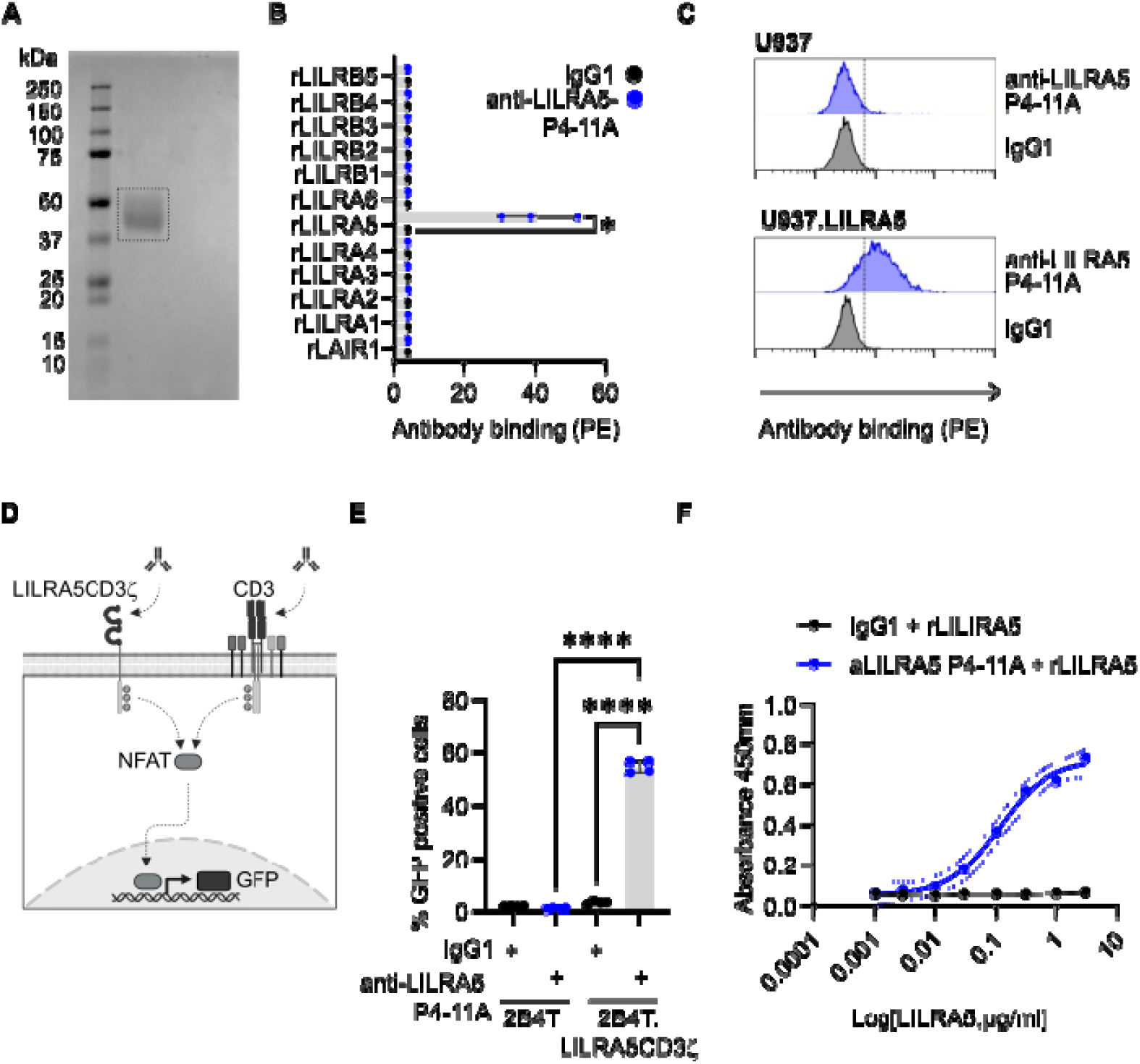
A highly specific anti-LILRA5 antibody with cross-linking capacity. **(A)** SDS-PAGE analysis of rLILRA5-His, used for immunisation of Balc/c mice. **(B)** Binding of anti-LILRA5 P4-11A mAb to magnetic beads coated with recombinant LILR or control proteins. mAb binding was detected using anti-IgG mAb and flow cytometric analysis. Mean ± SD of *n* = 3 independent experiments is shown. Paired *t*-test between anti-LILRA5 P4-11A and isotype control, where \**p* < 0.05. **(C)** U937 cells transfected with DNA vectors expressing LILRA5 protein, or control cells, were analysed for binding of anti-LILRA5 clone P4-11A mAb. mAb binding was detected using anti-IgG mAb and flow cytometric analysis. *n* = 3 from 3 independent experiments, one representative experiment is shown. **(D)** A schematic of the LILRA5CD3ζ reporter 2B4 T cell line, expressing a surface protein composed of extracellular and transmembrane LILRA5 domains fused to the cytoplasmic tail of CD3ζ. The cross-linking of CD3 or the fusion LILRA5CD3ζ protein induces phosphorylation of the ITAM (immunoreceptor tyrosine-based activation motif) domains in CD3ζ by Src kinases, and a signalling cascade that activates the NFAT (nuclear factor of activated T-cells) transcription factor. NFAT subsequently induces the expression of green fluorescent protein (GFP). **(E)** Cross-linking capacity of plate-bound anti-LILRA5 P4-11A mAb assessed using a LILRA5CD3ζ reporter cell line. GFP expression by LILRA5CD3ζ+ 2B4T cells or control 2B4T cells was assessed by flow cytometric analysis after incubation in wells containing plate-bound anti-LILRA5 P4-11A or isotype mAb. The % of GFP+ cells were quantified. Mean ± SD of *n* = 4 independent experiments are shown. One-way ANOVA, where \*\*\*\**p* < 0.0001. **(F)** Development of an ELISA for detecting sLILRA5. anti-LILRA5 P4-11A mAb or isotype control were used to capture rLILRA5-His. A titration curve (*n = 3*) of rLILRA5-His detected by anti-LILRA5 P4-11A mAb is shown.

As cross-linking is crucial for studying mAb-induced LILRA5 function, we tested whether anti-LILRA5 P4-11A mAb could cross-link and activate LILRA5 in a reporter cell model. We generated 2B4T NFAT-GFP reporter cells that express the LILRA5CD3ζ fusion protein at the cell surface (**Fig. 1D**). The engagement of CD3ζ in the control 2B4T NFAT-GFP cells and the LILRA5CD3ζ-expressing 2B4T NFAT-GFP induced GFP expression (**Supp. Fig. 2A**). Plate-bound rabbit polyclonal anti-LILRA5 antibodies induced the expression of GFP in LILRA5CD3ζ-expressing, but not control, 2B4T NFAT-GFP cells (**Supp. Fig. 2B**). This confirmed that cross-linking LILRA5CD3ζ chimaera induces GFP expression. Plate-bound anti-LILRA5 P4-11A mAb induced the expression of GFP in LILRA5CD3ζ-expressing, but not control, 2B4T NFAT-GFP cells (**Fig. 1E**). In contrast, incubation of either cell on plate-bound isotype IgG1 did not induce GFP expression. Therefore, anti-LILRA5 P4-11A mAb has agonistic properties for LILRA5 that can be utilised as a molecular tool to study LILRA5 signalling and functions.

Since soluble LILRA5 (sLILRA5) can be expressed by human immune cells (8,9), we investigated if anti-LILRA5 P4-11A could be used as a capture antibody in an ELISA assay. We used rLILRA5-His as a standard and rabbit anti-LILRA5 polyclonal antibodies for detection. The ELISA detected rLILRA5-His, but not rLAIR1-His as a control protein, in a concentration-dependent manner **(Fig. 1F)**. Based on these results, we conclude that the antiLLILRA5 P4-11A antibody only binds to human LILRA5, does not crossLreact with other LILRs and has cross-linking capacity.

### LILRA5 is expressed on the surface of human neutrophils and monocytes

The expression of *LILRA5* transcripts has been detected in primary human neutrophils and monocytes from healthy donors (9), but LILRA5 protein levels on neutrophils was never reported (10). We revisited the issue of transcript and protein expression, by analysing contemporary RNA-seq data from two independent studies and by using the anti-LILRA5 P4-11A mAb to detect protein expression with high-specificity. Analysis of RNA-seq data showed that *LILRA5* transcript abundance was comparable between neutrophils and monocyte populations, but at a lower level on dendritic cells (DCs) (**Fig. 2A**). These results indicate that human monocytes and human neutrophils express *LILRA5* transcripts at comparable levels. The binding of PE-conjugated anti-LILRA5 P4-11A to the surface of primary immune cells from *n* = 3 healthy donors detected expression of LILRA5 on the surface 95.58 ± 7.44 % of monocytes and 99.57 ± 0.47 % of neutrophils (**Fig. 2B** and **Supp. Fig. 3**). Analysis of the binding of anti-LILRA5 P4-11A to the surface of purified monocytes (anti-LILRA5 5.28 ± 3.05 vs. IgG1 1.17 ± 0.25) (**Fig. 2C** and **2D**) and purified neutrophils (anti-LILRA5 1.55 ± 0.26 vs. IgG1 1.24 ± 0.19) (**Fig. 2C** and **2E**) confirmed that both cell types express LILRA5. Notably, the surface LILRA5 expression was lower for neutrophils when compared to monocytes. Taken together, this reveals that naïve neutrophils and monocytes express comparable transcript levels of *LILRA5*, while monocytes express higher protein levels of LILRA5 on their cell surfaces.

**Figure 2:**
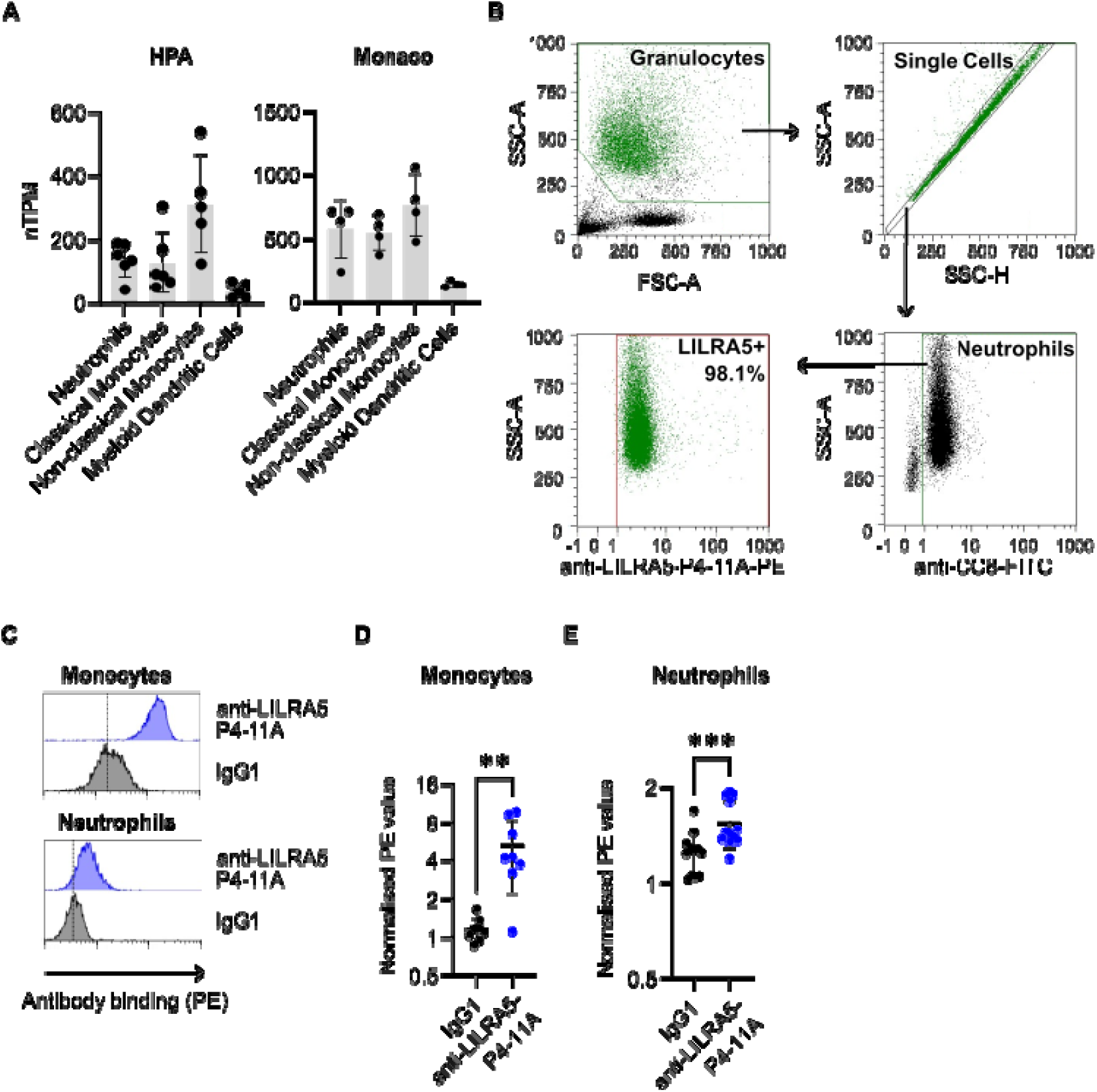
LILRA5 is expressed on human phagocytes. **(A)** The normalised transcripts per million (nTPM) from the HPA dataset and the Monaco dataset are shown. nTPM values give a quantification of the transcript abundance which is comparable across samples and genes. nTPM values for classical monocytes that have been characterised to express LILRA5 is shown as a control. Mean ± SD are shown. **(B)** Representative example showing the gating strategy used to identify granulocyres in human whole blood using anti-CEACAM8 (CC8; clone 6/40c) and anti-LILRA5 (clone P4-11A). A first gate was set on physical parameters of SSC-A vs. FSC-A, then on SSC-A and SSC-H to eliminate doublets, then granulocytes and monocytes were gated on CD14+ and CEACAM8+, then on CEACAM8+ events to identify granulocytes. **(C, D** and **E)** Expression of LILRA5 on human monocytes and neutrophils analysed by flow cytometry analysis. C shows a representative FACS staining indicating LILRA5 expression in neutrophils and monocytes from a healthy donor. D and E show quantification of the geometric mean of monocytes (*n* = 8 independent donors) and neutrophils (*n* = 12 independent donors) stained with anti-LILRA5 clone P4-11A or isotype control, respectively. Paired t-test, where \*\**p* < 0.01 and \*\*\**p* < 0.001.

### Respiratory burst can be stimulated through LILRA5

Next, we investigated whether LILRA5 cross-linking via anti-LILRA5 P4-11A could induce respiratory burst in neutrophils and monocytes. In these assays, we incubated human neutrophils or PBMCs in the presence of plate-bound anti-LILRA5 P4-11A mAb or isotype IgG1 control. We used anti-CD89 mAb as a positive control as it induces ROS production by cross-linking FcαR (CD89) and inducing a signal through FcRγ (14,15). Incubation of PBMCs from healthy donors with plate-bound anti-CD89 mAb or anti-LILRA5 P4-11A mAb, but not isotype IgG1 control, resulted in the ROS production (**Fig. 3A** and **3B**). As LILRA5 and FcαRI are not expressed on lymphocytes, ROS production is most likely elicited through the monocyte population in the PBMCs. Of note, LILRA5-induced ROS production was almost as potent as FcαR-induced ROS production. Likewise, neutrophils from healthy donors produced ROS in response to plate-bound anti-CD89 mAb and anti-LILRA5 P4-11A mAb, but not isotype IgG1 control (**Fig. 3C** and **3D**). This data indicates that LILRA5 can induce respiratory bursts in human monocytes and neutrophils.

**Figure 3:**
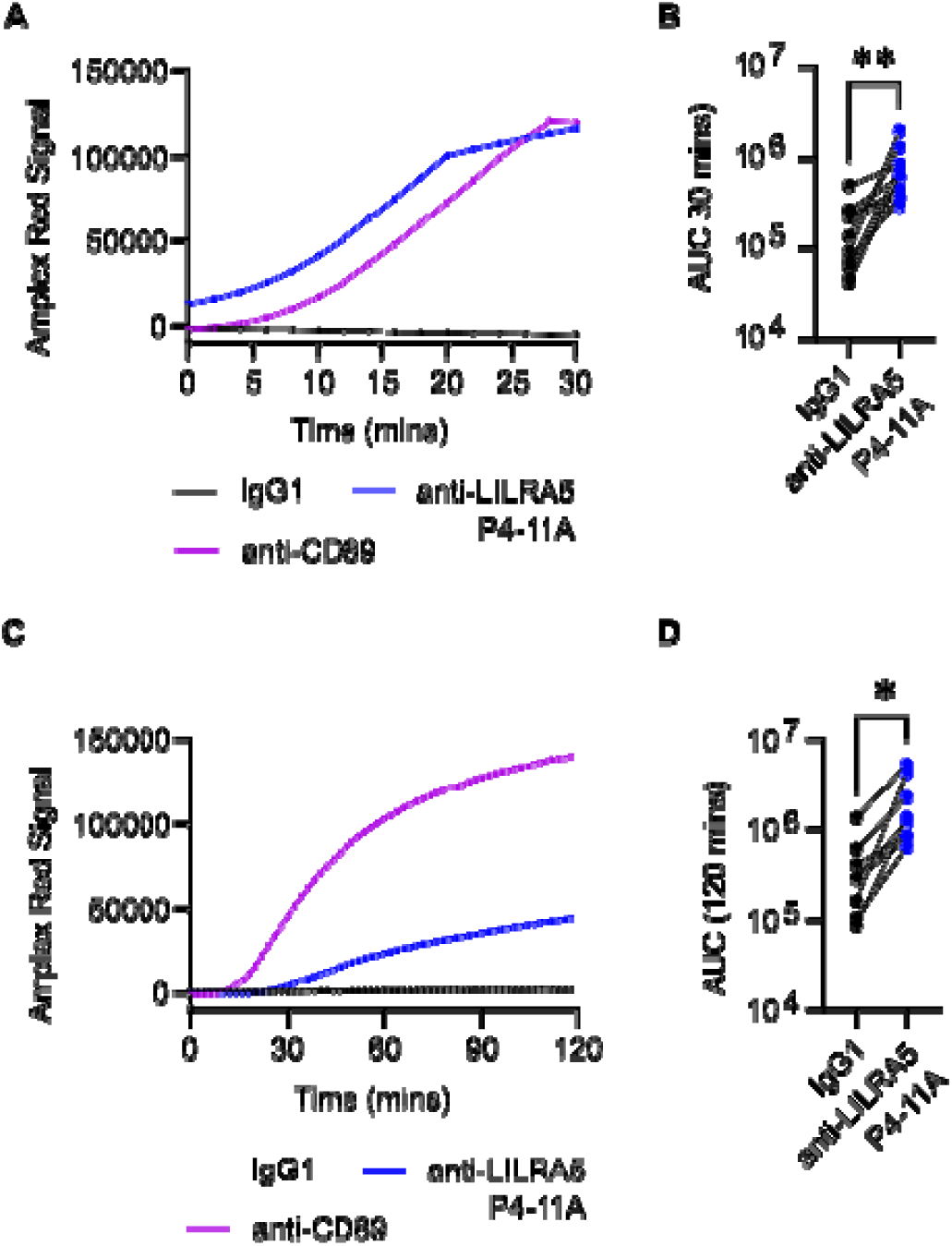
LILRA5 stimulates ROS production and in human phagocytes. **(A** and **B)** Stimulating LILRA5 on PBMCs induces reactive oxygen species (ROS) production, as measured using Amplex Red. A representative plot is shown in A. Mean ± SD of *n* = 7 independent donors is shown in B. A paired-sample *t*-test was used to compare Area Under the Curve (AUC) after stimulating PBMCs with anti-LILRA5 P4-11A or isotype IgG1 control, where \**p* < 0.05. **(C** and **D)** Stimulating LILRA5 on neutrophils induces ROS production, as measured using Amplex Red. A representative plot is shown in C. Mean ± SD of *n* = 8 independent donors is shown in D. A paired-sample *t*-test was used to compare AUC after stimulating neutrophils with anti-LILRA5 P4-11A or isotype IgG1 control, where \**p* < 0.05.

### *LILRA5* transcripts are elevated in systemic infection and sepsis

It was recently reported that *LILRA5* expression in monocytes is a biomarker of sepsis, a life-threatening organ dysfunction that occurs when the immune system overreacts to an infection (13,16). Therefore, we investigated the expression properties of *LILRA5* transcripts and LILRA5 protein in sepsis patients. We first assessed whether enhanced *LILRA5* expression was increased in whole blood of sepsis patients compared to healthy controls in additional datasets that were not analysed in previous studies. The elevated *LILRA5* expression in whole blood in sepsis patients was evident in several independent datasets **(Fig 4A** and **Supp Fig. 4A, 4B** and **4C)**. As previous analysis revealed that monocytes display enhanced *LILRA5* transcripts during sepsis (13,16), we questioned whether neutrophils as well as monocytes also display enhanced *LILRA5* expression in sepsis patients. We addressed this by analysing the Demaret *et al.* dataset (17), which revealed that *LILRA5* expression was enhanced in neutrophils purified from the peripheral blood of sepsis patients compared to healthy controls **(Fig. 4B).** Expression of *LILRA5* transcripts was also elevated in monocytes from sepsis patients compared to healthy controls in a previously unanalysed dataset **(Fig. 4C)**. Thus, *LILRA5* expression is increased in whole blood, neutrophils and monocytes during human sepsis.

**Figure 4:**
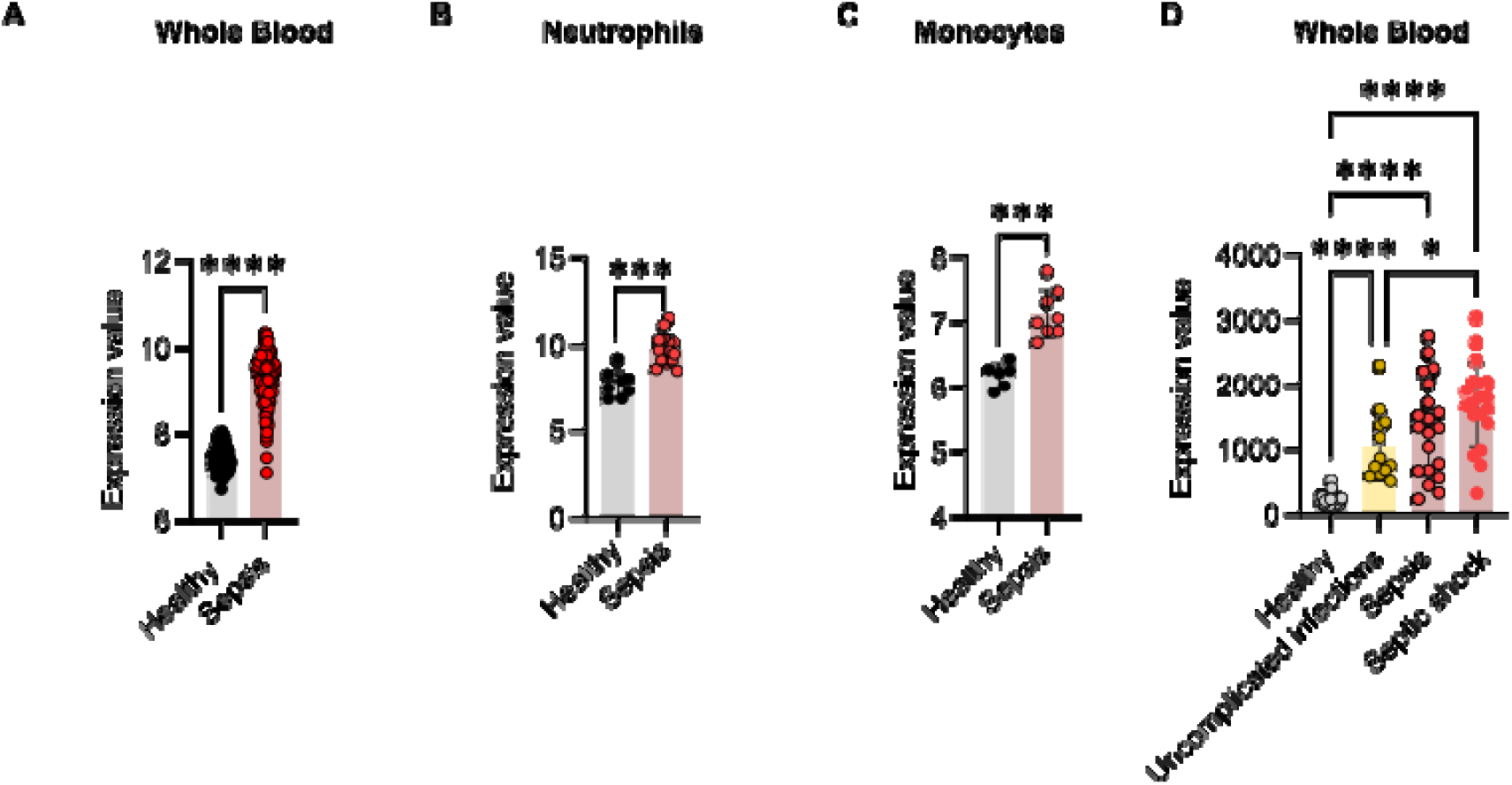
***LILRA5* expression in sepsis patients. (A)** *LILRA5* expression in whole blood of healthy donors (*n* = 83) or sepsis patients (*n* = 156) (GSE134364). Mean ± SD are shown. Statistics tested by *limma*, where adjusted \*\*\**p* < 0.001. **(B)** *LILRA5* expression on neutrophils from healthy donors (*n* = 8) or sepsis patients (*n* = 15) (GSE64457). Mean ± SD are shown. Statistics tested by *limma*, where adjusted \*\*\**p* < 0.001. **(C)** *LILRA5* expression in CD14+ monocytes from healthy donors (*n* = 6) or sepsis patients (*n* = 8) (GSE180387). Mean ± SD are shown. Statistics tested by *limma*, where adjusted \*\*\**p* < 0.001. **(D)** *LILRA5* expression in whole blood of healthy donors (*n* = 40), patients with diagnosed but uncomplicated systemic infection (n = 12), sepsis patients (*n* = 20) or septic shock patients (*n* = 19) (GSE154918). Mean ± SD are shown. Statistics tested by *limma*, where adjusted \*\*\**p* < 0.001.

To understand whether enhanced *LILRA5* expression is a marker of sepsis *per se*, it is important to establish whether patients with bacterial and viral infections that have not progressed to sepsis display modified *LILRA5* expression. First, we analysed the whole blood transcript dataset of Dix *et al.* from patients with diagnosed bacterial infections and controls (18). Here, *LILRA5* expression was increased in the whole blood of patients upon hospital presentation that were subsequently diagnosed with a *S. aureus* or *E. coli* invasive infection compared to healthy controls **(Supp Fig. 4D)**. Next, we analysed whole blood transcriptome data from healthy individuals, febrile children with confirmed bacterial infections and febrile children with confirmed viral infections from the Herberg *et al.* (19). This revealed that *LILRA5* expression is enhanced in individuals with bacterial infection or viral infection compared to healthy controls **(Supp Fig. 4E).** Finally, we analysed the dataset from Herwanto *et al.* to compare *LILRA5* expression in whole blood of healthy individuals, patients with diagnosed but uncomplicated infections, patients with sepsis and patients that progressed to septic shock (20). In this dataset, *LILRA5* transcripts in whole blood were significantly increased in patients with diagnosed but uncomplicated infections, in sepsis patients and in septic shock patients compared to healthy controls **(Fig. 4D)**. In sum, *LILRA5* expression is not significantly different between sepsis patients and patients with diagnosed but uncomplicated infections, suggestive that elevated *LILRA5* expression in whole blood is not a marker of sepsis *per se*.

### Surface LILRA5 expression does not increase upon immune challenge

Given the elevated *LILRA5* transcripts in whole blood during infectious and inflammatory diseases, we hypothesised that the expression of surface LILRA5 would be increased on immune cells under these conditions. A previous transcriptome analysis revealed that 4 hours of *ex vivo* infection of human blood with *E. coli*, but not *S. aureus,* induced a significant increase in expression of *LILRA5* **(Fig. 5A).** We assessed the binding of anti-LILRA5 P4-11A mAb to monocytes purified from human whole blood that had been infected with *E. coli* or *S. aureus*. Surprisingly, surface LILRA5 expression was not differentially regulated on monocytes after *ex vivo* infection with *E. coli* or with *S. aureus* **(Fig. 5B)**. Collectively, these results suggest that bacterial challenges induce elevated *LILRA5* expression in monocytes, but no altered surface LILRA5 expression on monocytes.

**Figure 5:**
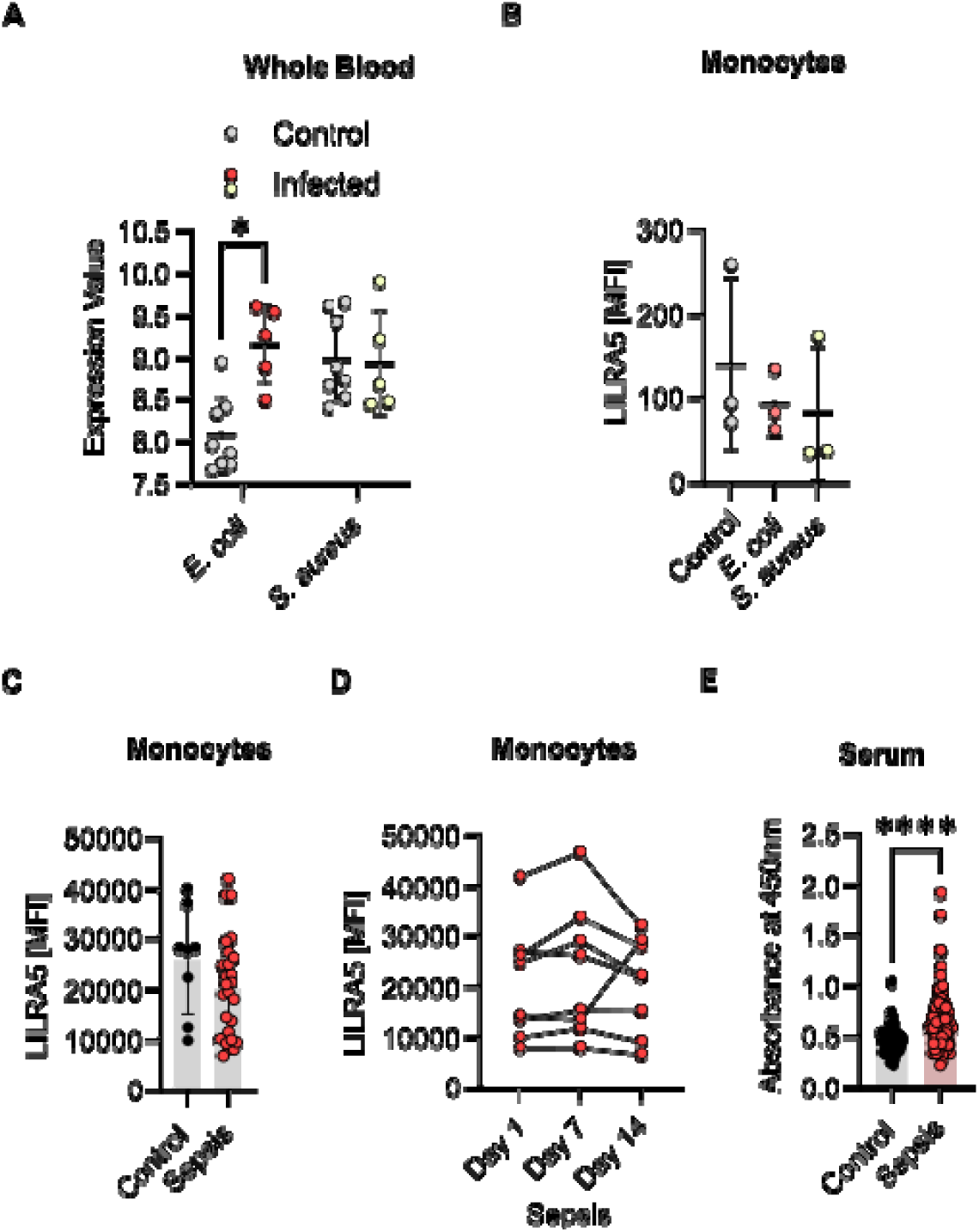
Surface LILRA5 expression does not increase during infection and inflammation. **(A)** *LILRA5* expression in whole blood from healthy donors after 4-hour *ex vivo* infection with *E. coli* (*n* = 5, control *n* = 8) or *S. aureus* (*n* = 5, control *n* = 8) (GSE65088). Mean ± SD are shown. Statistics tested by *limma* adjusted \*\*\**p* < 0.001. **(B)** Expression of LILRA5 on human monocytes after *ex vivo* infection of whole blood by *E. coli* or *S. aureus*, analysed by flow cytometry analysis. One-way ANOVA was used to compare the mean fluorescence intensity (MFI) from anti-LILRA5 P4-11A staining (after subtracting of MFI isotype IgG1) for infection vs control, where \*\**p* < 0.01 and \**p* < 0.05, is shown for *n* = 3 independent donors. **(C)** Surface LILRA5 expression on human monocytes from healthy donors (*n* = 8) and sepsis patients (*n* = 26), upon hospital admission, analysed by flow cytometry analysis. Each data point represents the MFI from anti-LILRA5 P4-11A stained cells after subtracting the MFI of isotype IgG1 stained cells. Mean ± SD are shown. **(D)** Surface LILRA5 expression on human monocytes from sepsis patients (*n* = 8) at day 1, day 7 and day 14 of hospital admission, analysed by flow cytometry analysis. Each data point represents the MFI from anti-LILRA5 P4-11A stained cells after subtracting the MFI of isotype IgG1 stained cells. **(E)** Comparison of sLILRA5 in serum from sepsis patients (*n* = 128) or healthy donors (*n* = 60). Mean ± SD are shown. Statistics were tested by student t-test, where \*\*\*\**p* < 0.0001.

To ascertain whether our *ex vivo* findings were representative of *in vivo* immune challenge, we examined the binding of anti-LILRA5 P4-11A mAb to PBMCs purified from sepsis patients and healthy controls. The sepsis patients had confirmed infections with Gram-negative bacteria (*n* = 12; 46 %), Gram-positive bacteria (*n =* 6; 23 %) or Candida (*n =* 3; 12 %), or unidentified aetiology (*n* = 4; 15 %). We specifically measured the LILRA5-signal on monocytes by co-staining the PBMCs with APC-conjugated anti-CD14, PE-conjugated HLA-DR and FITC-conjugated anti-LILRA5. Notably, there was no significant difference in surface LILRA5 expression levels on monocytes from sepsis patients at day 1 of diagnosis compared to healthy controls **(Fig. 5C)**. Furthermore, the surface LILRA5 expression on monocytes remained largely unchanged during the progression of sepsis in individual patients **(Fig. 5D)**. Therefore, *LILRA5* transcripts expression in innate immune cells is enhanced during immune challenge, but the surface LILRA5 expression remains unchanged.

### Soluble LILRA5 is elevated in serum from sepsis patients

Given our previous observations we queried how *LILRA5* transcripts could increase upon systemic infection and inflammation, but surface LILRA5 expression levels remained unchanged. As *LILRA5* can undergo alternative mRNA splicing to generate transcripts that encodes two different sLILRA5 isoforms (9), a switch in mRNA splicing could explain our previous results. Alternatively, our results could arise if there was no switch in mRNA splicing but surface LILRA5 was shed from cell surfaces. Under either scenario, it would be expected that sLILRA5 levels are elevated under immune challenge. To test this, we compared sLILRA5 in serum from sepsis patients and healthy donors using the ELISA assay and anti-LILRA5 P4-11A. The mean signal for sLILRA5 in serum from 128 patients with clinical defined sepsis was significantly higher than in serum from 60 healthy controls **(Fig. 5E)**. Taken together, this suggests that immune challenge increases sLILRA5 levels, indicative of splice changes or cell shedding.

### Immune challenge impacts LILRA5-induced ROS production

To investigate the functional consequences of altered *LILRA5* transcripts or surface LILRA5 expression, we challenged PBMCs with the potent immune stimulator LPS. LPS induces a significant increase in *LILRA5* transcript expression in monocytes **(Fig. 6A)**, and lead to a trend (*p* = 0.0683) for increased surface LILRA5 expression **(Fig. 6B)**. Since there was an increased surface LILRA5 expression on monocytes during LPS stimulation, we assessed the impact of this dynamic change on cellular function. As in our previous experiments, we measured the capacity of anti-LILRA5 P4-11A to induce ROS production when cells were pre-stimulated with LPS. Surprisingly, there was a significant reduction in LILRA5-dependent ROS production for cells that had been stimulated with LPS compared to untreated cells (**Fig. 6C** and **6D**). This indicates that LPS challenge increases *LILRA5* transcripts and expression of surface and soluble LILRA5, but that LILRA5 is less effective at inducing respiratory burst.

**Figure 6.**
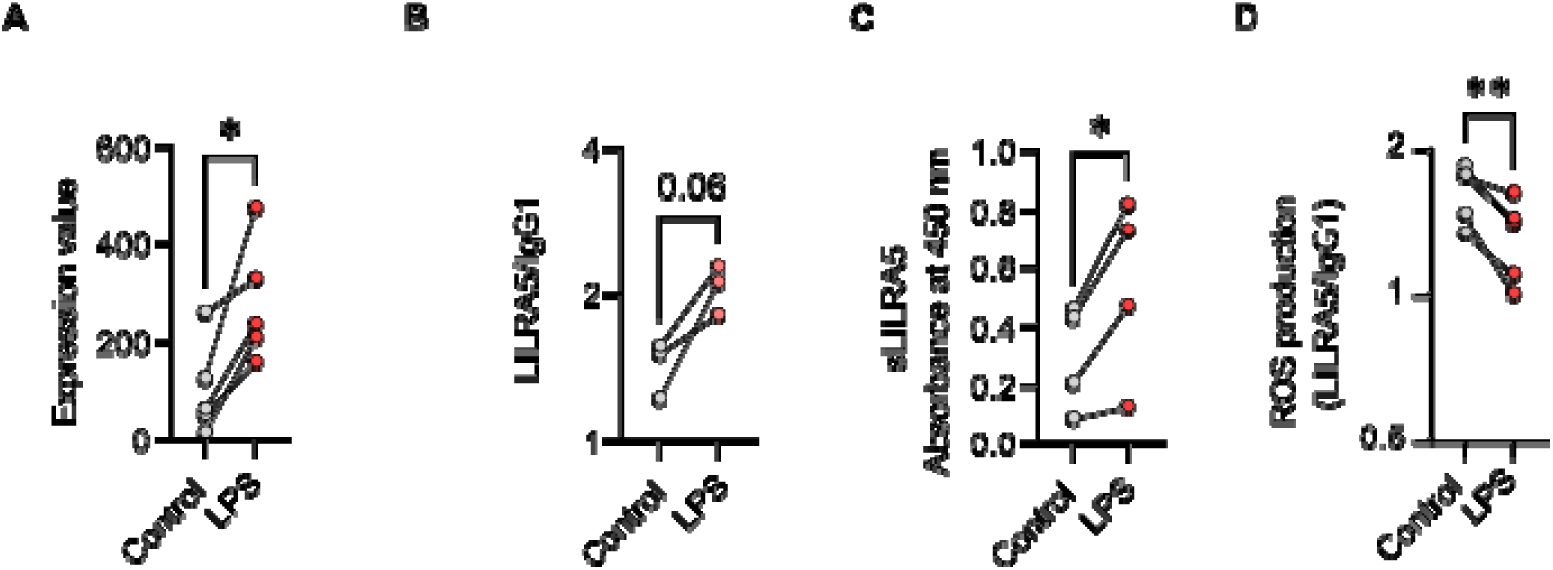
LPS activation reduces LILRA5-dependent ROS production. **(A)** *LILRA5* expression by monocytes purified from healthy donors that were cultured ± LPS for 18 hours (GSE147310). Data from *n* = 5 donors are shown. Statistics tested by *limma*, where adjusted \**p* < 0.05. **(B)** Surface LILRA5 expression on human monocytes purified from healthy donors (*n* = 3), analysed by flow cytometry analysis. Each data point represents the signal of anti-LILRA5 P4-11A relative to isotype IgG1 control. Statistics tested by paired t-test, where \**p* < 0.05. **(C)** sLILRA5 in culture supernatants from PBMCs purified from healthy donors (*n* = 4), after 18 hours culture ± LPS. Statistics were tested by paired t-test, where \**p* < 0.05. **(D)** Production of reactive oxygen species (ROS) by PBMCs in response to LILRA5 ± LPS stimulation, as measured using Amplex Red. ROS production was quantified as the area under the curve (AUC). The ROS production induced by anti-LILRA5 P4-11A relative to IgG1 was calculated. Data is shown from *n* = 5 independent donors. Paired *t*-test was relative ROS production induced by anti-LILRA5 P4-11A for control vs LPS-treated cells, where \*\**p* < 0.01.

## Discussion

Here, we demonstrate that human phagocytes express surface LILRA5 and that it promotes important effector functions of respiratory burst and degranulation. We show that LILRA5 is expressed on the surface of both circulating neutrophils and monocytes in healthy donors. Since oxidative burst is induced by immune receptors that act to sense non-self and that signal through ITAMs (21), our data imply that LILRA5 has a role in mounting innate immune responses. Enhanced ROS production, mediated through a LILRA5-dependent mechanism during infection or immune challenge, would contribute to the killing of microbes within the phagolysosome following successful phagocytosis. Indeed, deletion of LILRA5 in mice led to enhanced progression of *P. aeruginosa*-induced inflammation and keratitis (12). Additionally, the increased release of antimicrobial factors, such as proteases and peptides, into the local environment following LILRA5-triggered exocytosis of granules, would further promote microbial killing.

The level of activating receptor expression on an immune cell is an important determinant of its capacity to induce cellular activation. Notably, a substantially higher expression of surface LILRA5 expression is present on circulating monocytes compared to neutrophils in healthy individuals, even though there is a comparable level of *LILRA5* transcripts. This differential expression suggests that LILRA5 may have distinct roles on monocytes and neutrophils. Interestingly, even with the lower surface expression of LILRA5 on neutrophils, our assays demonstrate that ROS production can still be effectively induced via LILRA5 cross-linking. This indicates that LILRA5, although less abundant on neutrophils, is functionally competent and capable of triggering significant immune responses. The higher expression of LILRA5 on the surface of monocytes might reflect a more prominent role in sustained immune activation and regulation, whereas the presence on neutrophils, albeit at lower levels, underscores its importance in rapid, acute responses to infection

The immune system initiates a swift immune response to detect non-self in order to protect the host. The upregulation of *LILRA5* expression at transcript levels in whole blood, monocytes and neutrophils in response to infection or immune challenge, suggests that it has a role in sensing or mounting responses against microbes and/or non-self. In line, it has previously been reported that expression of mRNA encoding membrane-anchored LILRA5 in monocytes is dynamic and increases upon exposure to cytokines (8). Given this, we hypothesised that the increased *LILRA5* expression upon immune challenge would be correlated with an increase in surface LILRA5 expression on immune cells. Surprisingly, our study revealed that surface LILRA5 expression does not change on the surface of monocytes during infection of bacterial pathogens *ex vivo*. Additionally, monocytes from sepsis patients displayed no change in surface LILRA5 expression compared to healthy controls. This could be explained if the increased *LILRA5* transcripts encode sLILRA5 but not membrane-bound LILRA5. *sLILRA5* expression is reported in multiple immune cell types and varies between individuals (22). In line, sLILRA5 was detected in sera of sepsis patients in our study, and in the synovial fluid of rheumatoid arthritis patients and in sera of patients with liver steatosis (8,23). Our studies did identify a trend for increased expression of surface LILRA5 on monocytes after their stimulation with LPS. These results suggest that surface LILRA5 expression is dynamic, changing in response to different stimuli and/or over time. Further investigation is needed to elucidate the precise mechanisms governing both surface and soluble LILRA5 expression and their functional implications in immune responses.

The function of an activating receptor is dependent on the sensing of a ligand, but LILRA5 has no defined ligand (7,10,11). Most activating receptors on phagocytes sense PAMPs, DAMPs, antibody-opsonised microbes or complement-opsonised microbes (21,24). However, LILRA2, a receptor in the same family, can sense microbially-cleaved antibodies to trigger antibacterial effector functions (25). This raises the prospect that LILRA5 could also sense infection by detecting modified self ligands. sLILRA5 may act to regulate membrane-anchored LILRA5 functions by competing for the same ligand. Thus, the identification of LILRA5 ligands is important to understand its biological and evolutionary properties.

Enhanced *LILRA5* expression in whole blood or monocytes has been reported as a sepsis biomarker in patient cohorts (12,13). However, *LILRA5* transcripts are also increased in patients with diagnosed bloodstream infections. Furthermore, *LILRA5* transcript levels in whole blood do not significantly differ between patients with diagnosed but uncomplicated infection and patients with sepsis. Since levels of *LILRA5* transcripts cannot differentiate these two patient groups, this suggests that *LILRA5* is not a biomarker of sepsis *per se*. Our analysis of sLILRA5 in serum revealed elevated levels in sepsis patients compared to healthy controls. Additional quantitative studies are required to test the robustness of this finding, and to ascertain whether quantifying sLILRA5 in sera is a putative biomarker of sepsis.

In conclusion, this study was underpinned by the development of a novel LILRA5-targeting mAb, anti-LILRA5 clone P4-11A, that detects LILRA5 with high specificity and possesses agonistic properties. LILRA5 is expressed by human phagocytes and promotes ROS production. Immune challenge can induce dynamic increases in *LILRA5* transcript levels, but unchanged or increased surface LILRA5 expression. This regulates LILRA5-dependent ROS production. Future studies are required to identify LILRA5 ligands and to further elucidate the regulatory mechanisms and functions of membrane-bound LILRA5 and sLILRA5.

## Materials & Methods

### Ethical Approval

Human blood was obtained from healthy donors, approved by the Regional Ethics Committee and Imperial College Healthcare NHS Trust Tissue Bank (Regional Ethics Committee approval no. 17/WA/0161, Imperial College Healthcare Tissue Bank Human Tissue Authority licence no. 12275, and Imperial College Research Ethics Committee no. 19IC5166). The study was approved by the local ethics committee of the Friedrich-Schiller-University Jena for samples from sepsis patients (ID:2019-1306_2-Material) and healthy donors (ID:2022-2847-Material). All samples were collected after receiving signed informed consent from all participants. Serum sepsis samples were approved by the Ethics Committee of Sumy State University (protocol number 12, approval date December 11th 2021).

### Expression and purification of rLILR-His

EXPI293F cells were cultured in EXPI media at 37°C with 5% CO_2_, as previously described (15,26). The expression of rLILRB1-His, rLILRB3-His and rLILRA6-His have been previously described (15). To construct expression vectors for LILRA5, the remaining LILR and LAIR-1, the signal peptide and extracellular domains of LILRs were amplified from cDNA vectors (**Table 1**) and inserted into pcDNA3.4 vectors. The forward primers contained a pcDNA3.4 backbone overhang, a *Bam*HI restriction site, Kozak sequence and a gene-specific region. The reverse primers contained pcDNA3.4 backbone, a *Not*I restriction site, a stop codon, a 6xHis tag and a gene-specific region. PCR amplification was performed using Phusion High-Fidelity Taq Polymerase (Thermo Fisher Scientific) and thermocycling as follows: 1 cycle (98°C for 2 min), 35 cycles (98°C 15 s, 62°C 30 s, and 72°C 30 s), and 1 cycle (72°C 10 min). Recombinant His-tagged proteins were purified from culture supernatants by affinity chromatography (ÄKTA pure; GE Healthcare Life Sciences) using HIStrap nickel columns (GE Healthcare Life Sciences), and elutions were dialysed into 50 mM Tris 300 mM NaCl (pH 8) and stored at -80°C.

**Table 1:**
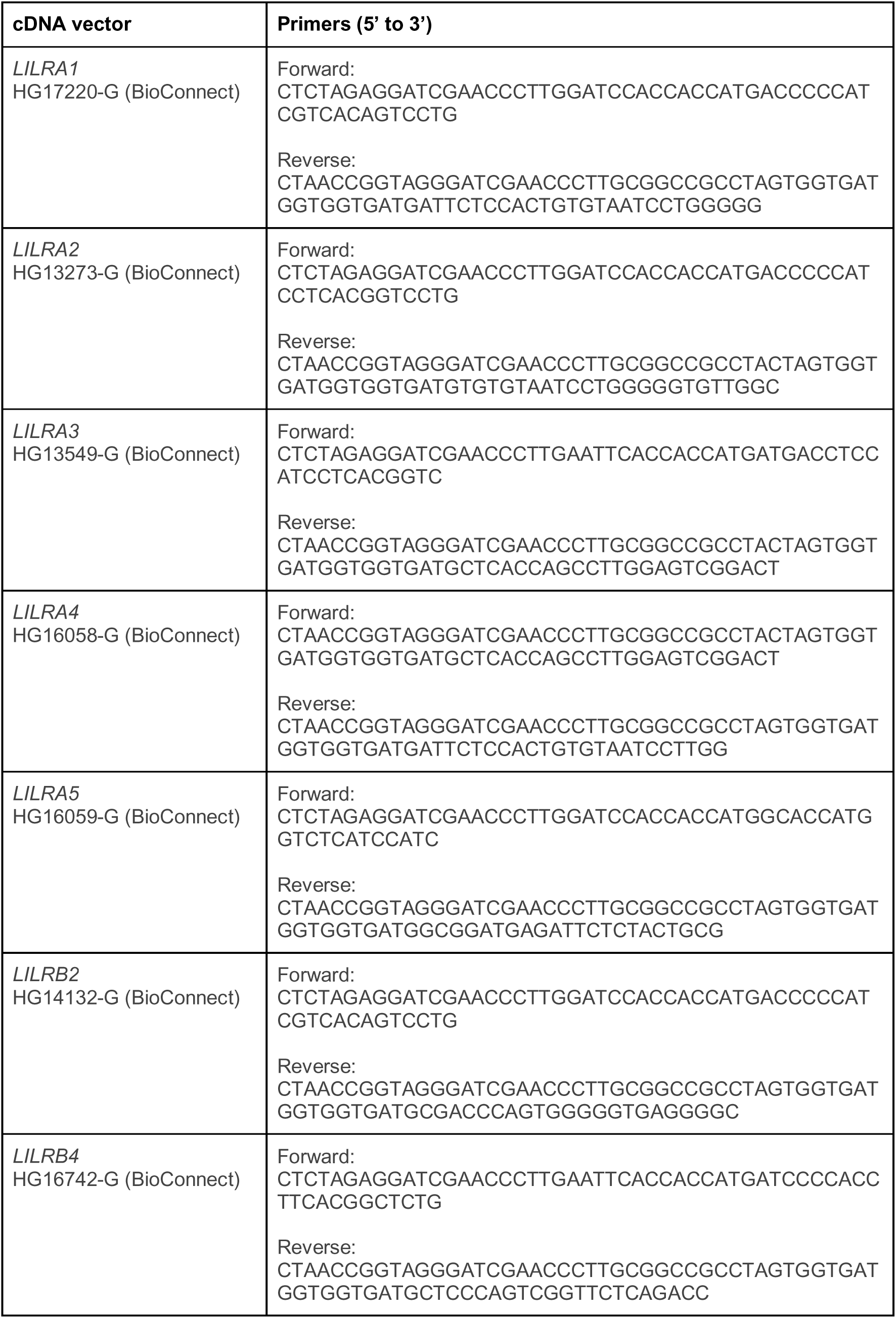
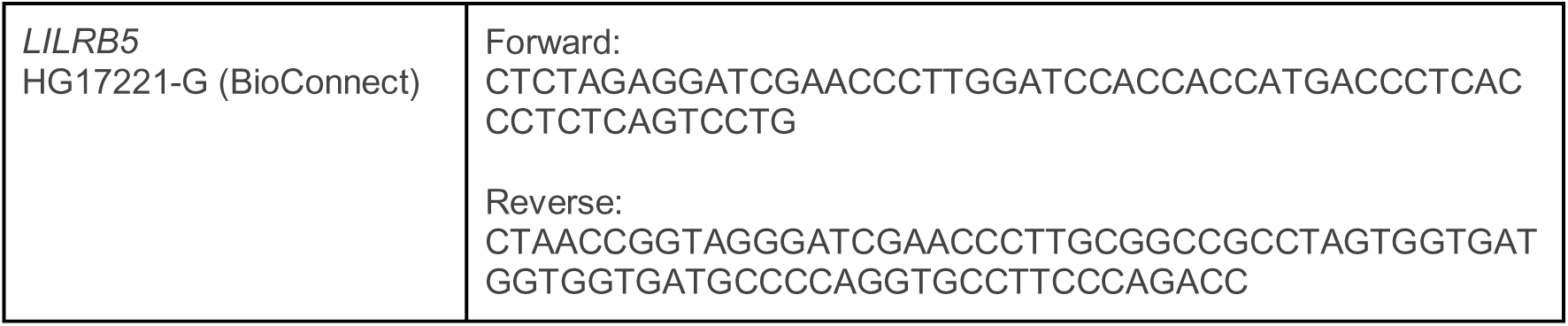
cDNA vectors and amplification primers.

#### Generation of anti-LILRA5 mAb

Generation of LILRA5-specific mAb was performed using the hybridoma technique of Köhler and Milstein (27). A Balb/c mouse was immunised with rLILRA5-His (David’s Biotechnologie, Germany). Spleen cells from Balb/c mice were recovered and fused to myeloma NS1/0 cells. The soluble IgG expressed by individual hybridoma clones were tested for binding rLILRA5-His by ELISA allowing the identification of clone P4-11A. The mAb was produced in a serum- and azide-free medium to ensure purity and minimise potential contaminants.

#### Binding of mAb to Dynabeads or cells

Dynabeads His-Tag Isolation and Pulldown (Thermo Fisher Scientific) were incubated with C-terminal His-tagged rLILR or rLAIR-1 proteins following manufacturer protocols to generated coated-DBs. DBs were incubated with 5 µg/ml of mAb in buffer (PBS + 0.5% BSA for DB) for 30 mins at 4°C. After washing, DB were incubated with 50 µl of PE-conjugated goat anti-mouse IgG secondary antibody (1:300; Invitrogen) under the same conditions. The fluorescence of DBs was measured by flow cytometry and analysed based on the FSC and SSC parameters.

#### Leukocyte immunophenotyping by flow cytometry

Hetasep (STEMCELL Technologies, Vancouver, Canada) was diluted in whole blood (1:7 v/v) and incubated for 30 min at room temperature. The supernatant was collected and washed with culture medium (RPMI1640 with stable glutamine (Bio&Sell, Feucht, Germany), 100 U/ml Penicillin, 100 μg/ml Streptomycin (both Gibco/ThermoFisher, Waltham, Massachusetts)) supplemented with 2% FCS (Biochrom; v/v). Platelets were removed by centrifugation at 200 g for 10 min without brake. Residual red blood cells were lysed with Red Blood Cell Lysis Solution (Miltenyi Biotec, Bergisch Gladbach, Germany) according to the manufacturer’s instructions. Isolated leukocytes were incubated with human TruStain FcX (Biolegend, San Diego, CA, USA) and stained with 10 µg anti-CD14-APC (Immunotools, Friesoythe, Germany), anti-CEACAM8 (clone 6/40c), and anti-LILRA5-11A (in house labeled with PE) each. Staining with the corresponding isotype control antibodies (mIgG-FITC, mIgG-PE, mIgG-APC; Immunotools, Friesoythe, Germany) was performed to determine the threshold for specific staining. Data were acquired using MACSQuant® Analyzer 10 (Miltenyi Biotec, Bergisch Gladbach, Germany) and analysed using MACSQuantify™ Software 2.13 (Miltenyi Biotec, Bergisch Gladbach, Germany). Monocytes were gated as CD14+ cells and neutrophils as CEACAM8+ cells. The median fluorescence intensity (MFI) of LILRA5 or isotype control was determined. Individual expression of LILRA5 was calculated as [MFI LILRA5 - MFI Isotype control].

### Isolation of primary human neutrophils and PBMCs

Neutrophils and PBMCs were isolated from whole blood by Ficoll/Histopaque centrifugation and resuspended in RPMI1640 supplemented with 5% FCS, as previously described (15). Neutrophils were isolated with >98% purity and 99% viability.

#### Binding of mAb to cells or primary immune cells

45 µl of 5 x 10L cells/ml of cell lines or purified primary immune cells were incubated with 5 µg/ml of mAb in buffer (PBS + 0.5% BSA for DB, or RPMI1640 supplemented with 10% FCS for cells) for 30 min at 4°C. After washing, cells were incubated with 50 µl of PE-conjugated goat anti-mouse IgG secondary antibody (1:300; Invitrogen) under the same conditions. The fluorescence of cell lines or purified primary immune cells was measured by flow cytometry and analysed based on the FSC and SSC parameters.

#### ELISA for detection of sLILRA5

Wells of 96-well plates were coated overnight with 50 µl of 2 µg/ml anti-LILRA5 clone P4-11A mAb in PBS at 4°C. After washing (3 times with PBS + 0.05% Tween-20), wells were blocked with 100 µl PBS + 5% BSA for an hour at room temperature. After three washes with PBS + 0.05% Tween-20, wells were incubated with 100 µl of rLILRA5-His or rLAIR1-His (at varying concentrations) in buffer or heat-inactivated human serum samples for 1 hour at 4°C. After washing (3 times with PBS + 0.05% Tween-20), 50 µl of rabbit anti-LILRA5 pAb (1:2000; Sino Biological) was added to the wells and incubated as above. For the detection of the rabbit anti-LILRA5 pAb, wells were incubated with 50 µl of horseradish peroxidase (HRP)-conjugated goat anti-rabbit-IgG antibody (1:10,000; A16096; Life Technologies) under the same condition. After washing (3 times with PBS + 0.05% Tween-20), the assay was developed with 100µl 1xTMB substrate and 100µl Stop solution for TMB substrate (Invitrogen).

#### Generation of LILRA5-expressing U937 cell line

U937 cell lines were cultured in RPMI1640 supplemented with 10% FCS and penicillin/streptomycin at 37°C with 5% CO_2_. To construct a LILRA5-expressing U937 cell line, a DNA fragment containing the coding domain sequence of LILRA5 was amplified from a LILRA5 cDNA vector (HG16059-G, BioConnect) using primers ^5’^GAGCTAGCAGTATTAATTAACCACCATGGCACCATGGTCTCATCCATC^3’^ and ^5’^GTACCGGTTAGGATGCATGCTCACCTTCCAGCTGCAGCTTGGG^3’^. PCR amplification was performed using Phusion High-Fidelity Taq Polymerase (Thermo Fisher Scientific) and thermocycling as follows: 1 cycle (98°C for 2 min), 35 cycles (98°C 15 s, 60°C 30 s, and 72°C 30 s), and 1 cycle (72°C 10 min). The purified amplicon was ligated into a dual promoter lentiviral vector (BIC-PGK-Zeo-T2a-mAmetrine; RP172 derived from no.2025.pCCLsin.PPT.pA.CTE.4⋅-scrT.eGFP.mCMV.hPGK.NG-FR.pre, as previously described (15) via Gibson reaction using a parental vector digested with *Pac*I and *Sph*I. Lentiviral particles were created in HEK293T cells and transfected into U937 cells, as previously described (15,28). Transfected cells were selected by culture in media supplemented with 400 μg/mL zeocin.

#### Generation of LILRA5CD3**ζ** reporter cell line and GFP induction

2B4 NFAT-GFP T cell reporter cells contain an NFAT-GFP fusion construct containing three tandem NFAT-binding sites fused to enhanced GFP cDNA (28). 2B4 T cell lines were cultured in RPMI1640 supplemented with 10% FCS and pen/strep at 37°C with 5% CO_2_. A LILRA5CD3ζ reporter cell line was generated, as previously described (15). In brief, a DNA fragment containing the coding domain sequence of the LILRA5 extracellular and transmembrane domains and the CD3ζ cytoplasmic tail was synthesised by Integrated DNA Technologies. The fragment was ligated into the RP172 vector, as described above. After selection and expansion, the resultant cell cultures were tested for LILRA5 expression using anti-LILRA5 pAb (16059-RP01; Sino Biological) and flow cytometry analysis. To induce GFP expression, TC-treated 96-well plates were coated at 4°C overnight with 40 μl of 5 μg/ml Hamster anti-CD3ζ clone 145-2C11 (BD Biosciences), anti-LILRA5 pAb (16059-RP01; Sino Biological), anti-LILRA5 P4-11A mAb, or IgG isotype control mAbs diluted in PBS. After washing, 96-well plates were seeded with two hundred microliters of 2B4T cells at a density of 2.5 × 10^5^ cells/ml in RPMI 1640 containing 10% FCS, 100 μg/ml penicillin, and 100 μg/ml streptomycin. After centrifugation at 200 rpm for 3 mins, cells were cultured at 37°C with 5% CO_2_ for 18 hours. The cells were then resuspended in PBS and fluorescence was measured by flow cytometry analysis.

#### Reactive oxygen species (ROS) production upon LILRA5 cross-linking

To quantify ROS production, wells of a white 96-well plate was coated overnight at 4°C with 50 µl of 5 µg/ml anti-LILRA5 clone P4-11A mAb, anti-CD89 (MCA1824; Bio Rad) or IgG1 Isotype control (0102-01; Southern Biotech) mAbs diluted in NaHCOℒ buffer, pH8.6. The plate was then washed twice with PBS. Neutrophils and PBMCs were washed with ice-cold reaction buffer (20 mM HEPES, 140 mM NaCl, 1 mM CaClℒ, 5 mM Glucose, pH7.4) three times and resuspended to 1 x 10L cells/ml. To each well, 100 µl of 1 x 10L cells and 100 µl of pre-warmed reaction buffer containing 20 mM Amplex Red (Life Technologies) were added. For the assays performed with PBMCs, the reaction buffer was additionally supplemented with 10 µl of horseradish peroxidase (1 unit/ml; P8125-5KU; Sigma Aldrich). Fluorescence was measured by a CLARIOstar microplate reader with detection at 2-min intervals for 2 hours at 37°C. Area under the curve (AUC) for each stimulus was calculated after subtraction of background/PBS ROS production. In specific experiments, cells were stimulated with 5 µg/ml LPS from *Escherichia coli* O111:B4 (Sigma–Aldrich) for one hour at 37°C with 5% CO_2_.

#### Activation of neutrophils and monocytes by LILRA5 cross-linking and/or LPS

Wells of a white 96-well plate were coated overnight at 4°C with 50 µl of 5 µg/ml anti-LILRA5 clone P4-11A mAb or IgG1 Isotype control (0102-01; Southern Biotech) mAbs diluted in PBS. The plate was then washed twice with PBS. Neutrophils and PBMCs were resuspended to 1 x 10L cells/ml in a solution made up of 80% of RPMI 1640 + penicillin/streptomycin and 20% of plasma from the same donor. To each well, 200 µl of 1 x 10L cells/ml was added and cultured for 24 hours at 37°C with 5% CO_2_. In certain assays, neutrophils and PBMCs were stimulated with 1 µg/ml or 10 ng/ml LPS from *Escherichia coli* O111:B4 (Sigma–Aldrich) for 24 hours at 37°C with 5% COL, respectively. Wells coated without antibodies and/or without LPS were used as a control. Neutrophils were collected and stained with anti-CD66b-FITC (1:50; BD Biosciences) for 30 mins at 4°C. In parallel, PBMCs were stained with 3μg/ml anti-CD14-FITC (367116, Biolegend) under the same conditions. After washing, the fluorescence of cells was measured by flow cytometry and analysed based on the forward- and side-scatter plots.

#### *LILRA5* expression profiling in transcriptome studies

The expression of *LILRA5* was assessed in retrieved data from previous array profiling data available on the Gene Expression Omnibus (GEO–NCBI) using GEO2R (NCBI). Datasets analysed were as follows:-1) to assess *LILRA5* expression in whole blood of sepsis patients and controls = GSE134364 from (29), GSE137342 from Mukhopadhyay, GSE185263 from (30), 2) to assess *LILRA5* expression in neutrophils purified from sepsis patients and controls = GSE64457 from (17), 3) to assess *LILRA5* expression in monocytes purified from sepsis patients and controls = GSE180387 from (31), 4) to assess *LILRA5* expression in whole blood of sepsis patients, septic shock patients, patients with diagnosed but uncomplicated infections and controls GSE154918 from (20), 5) to assess *LILRA5* expression in whole blood of patients with diagnosed bacterial or viral infections = GSE72810 from (19) and GSE33341 from (32), 6) to assess LILRA5 expression in whole blood after ex vivo infection with *E. coli* or *S. aureus* = GSE65088 from (18), 7) to assess LILRA5 expression in monocytes after culture for 18 hours ± LPS = GSE147310 from (33)

#### LPS-stimulation of leukocytes and quantification of surface LILRA5 and sLILRA5

Hetasep (STEMCELL Technologies, Vancouver, Canada) was diluted in whole blood (1:7 v/v) and incubated for 30 min at room temperature. The supernatant was collected and washed with culture medium (RPMI1640 with stable glutamine (Bio&Sell, Feucht, Germany), 100 U/ml Penicillin, 100 μg/ml Streptomycin (both Gibco/ThermoFisher, Waltham, Massachusetts)) supplemented with 2% FCS (Biochrom; v/v). Platelets were removed by centrifugation at 200 g for 10 min without brake. Residual red blood cells were lysed with Red Blood Cell Lysis Solution (Miltenyi Biotec, Bergisch Gladbach) according to the manufacturer’s instructions. Cultures were set up in culture medium supplemented with 4% FCS at 4x10^5^ cells/well/200 µl in 96-well ultra-low attachment culture plates (Corning, New York #3474) at 37°C with 5% CO_2_. After 60 min, cells were stimulated with 50 ng/ml *E.coli* 026:B6 LPS (Sigma, St. Louis, Missouri) for further 18 h. Leukocytes were incubated with human TruStain FcX (Biolegend, San Diego, CA) and stained with anti-CD14-APC (Immunotools, Friesoythe), anti-HLADR-PE (Invitrogen/ ThermoFisher Waltham, Massachusetts), and anti-LILRA5-11A (in house labeled with FITC) as described previously

(34). Staining with the corresponding isotype control antibodies was performed to determine the threshold for specific staining. The threshold was set at 3% false positive cells. Data were acquired using CytoFLEX S (Beckman Coulter, Krefeld, Germany) and analysed using CytExpert (Beckman Coulter). Monocytes were gated as CD14+HLA-DR+ cells and the median fluorescence intensity (MFI) of LILRA5 or isotype control was determined. Individual expression of LILRA5 was calculated as [MFI LILRA5 - MFI Isotype control].

#### Whole blood infection and measurement of surface LILRA5 expression on monocytes

To stimulate whole blood, 1 x 10L CFU of *E. coli* MG1655 or *S. aureus* JE2 were inoculated into 10 mL of whole blood for a 2-hour incubation at 37°C. PBMCs were then isolated using the standard protocol described previously. Cells (45 µL of 5 x 10L cells/mL) were incubated with 3 µg/mL IgG1 Isotype control (0102-01; Southern Biotech) or anti-LILRA5 P4-11A mAb in RPMI 1640 supplemented with 10% FCS for 30 minutes at 4°C. After washing with 1x PBS, cells were incubated with 50 µL of PE-conjugated goat anti-mouse IgG secondary antibody (1:300; Invitrogen) under the same conditions. The fluorescence of cells was measured by flow cytometry and analysed based on the forward- and side-scatter plots.

#### Analysis of monocytes and serum from sepsis patients

Sepsis was diagnosed according to the Sepsis-3 criteria (35). In total, *n* = 26 patients (16 males and 10 females) with sepsis were enrolled at the Department of Anesthesiology and Intensive Care Medicine, University Hospital Jena, Germany. The median (IQ range) age of the patients was 65 (53–72). The source of infection was Gram-negative bacteria (*n* =12; 46 %), Gram-positive bacteria (*n =* 6; 23 %), Candida (*n =* 3; 12 %), or unidentified (4; 15 %). Heparinised blood was drawn within 24 h after admission to the intensive care unit (ICU). If the patient remained on the ICU blood was collected up to two more times on day 7 and day 14 after admission. Whole blood from healthy donors (*n* = 9) with median age (IQ range) 52 (30–57) was used as control. The study was approved by the local ethic committee of the Friedrich-Schiller University Jena for samples from patients (ID:2019-1306_2-Material) and healthy donors (ID:2022-2847-Material) and of the University Hospital Essen (for samples from healthy donors). Informed consent was obtained from each patient or the respective legal representative and from healthy donors.

## Abbreviations

DAMP: Damage-associated molecular pattern
DB: Dynabeads
FcαR: Fc alpha receptor
FcRγ: Fc receptor gamma-chain
FCS: Foetal calf serum
FITC: Fluorescein Isothiocyanate
GFP: Green fluorescent protein
IFN-γ: Interferon gamma
IgG1: Immunoglobulin G1
IL-1β: Interleukin-1 beta
IL-6: Interleukin 6
ITAM: Immunoreceptor tyrosine-based activation motif
LAIR1: Leukocyte associated immunoglobulin–like receptor 1
LILRA2: Leukocyte Immunoglobulin like receptor A2
LILRA5: Leukocyte Immunoglobulin like receptor A5
LPS: Lipopolysaccharide
mAb: Monoclonal antibody
NFAT: Nuclear factor of activated T cells
PAMP: Pathogen-associated molecular pattern
PBMCs: Peripheral blood mononuclear cells
PE: Phycoerythrin
ROS: Reactive oxygen species
TNF-α: Tumour necrosis factor alpha

---

**Supplementary Figure 1:**
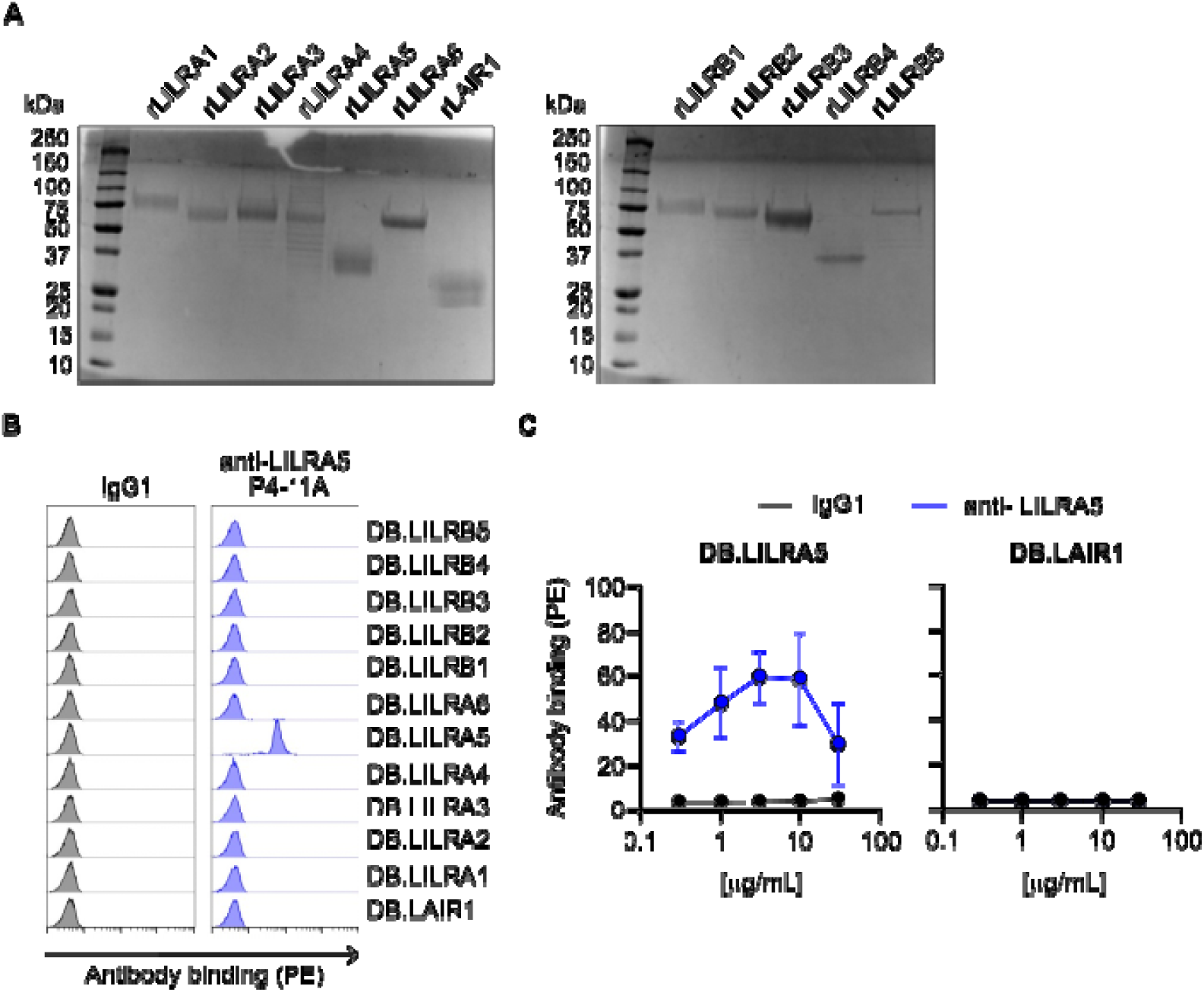
anti-LILRA5 P4-11A mAb binds to LILRA5 with high-specificity. **(A)** SDS-PAGE analysis of recombinant (r) proteins, used for assessing the specificity of anti-LILRA5 P4-11A mAb. **(B)** Specificity of anti-LILRA5 P4-11A antibody for LILRA5. Magnetic beads coated with rLILR or control proteins and analysed for binding of anti-LILRA5 P4-11A mAb. mAb binding was detected using anti-IgG mAb and flow cytometric analysis. *n* = 3 from 3 independent experiments, one representative experiment is shown. **(C)** Concentration-dependent binding of anti-LILRA5 P4-11A mAb to rLILRA5-coated dynabeads. mAb binding was detected using anti-IgG mAb and flow cytometric analysis. *n* = 3 from 3 independent experiments.

**Supplementary Figure 2:**
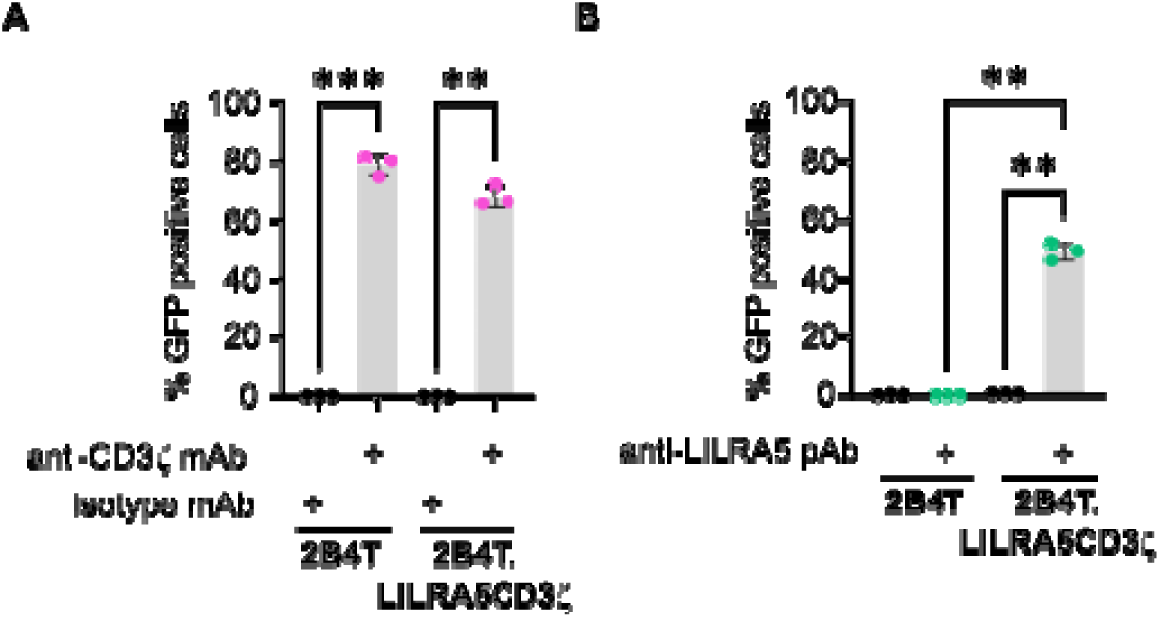
Validation of LILRA5. **(A)** Cross-linking capacity of plate-bound anti-CD3ζ mAb to induce GFP expression in the LILRA5CD3ζ reporter 2B4T cell line and control 2B4T cell line. GFP expression by LILRA5CD3ζ+ 2B4T cells or control 2B4T cells was assessed by flow cytometric analysis after incubation in wells containing plate-bound anti-CD3ζ or isotype mAb. The % of GFP+ cells was quantified. Mean ± SD of *n* = 3 independent experiments are shown. One-way ANOVA, where \*\*\**p* < 0.001, \*\**p* < 0.01. **(B)** Cross-linking capacity of plate-bound anti-LILRA5 pAb to induce GFP expression in the LILRA5CD3ζ reporter 2B4T cell line and control 2B4T cell line. GFP expression by LILRA5CD3ζ+ 2B4T cells or control 2B4T cells was assessed by flow cytometric analysis after incubation in wells containing plate-bound anti-LILRA5 or buffer control. The % of GFP+ cells were quantified. Mean ± SD of *n* = 3 independent experiments are shown. One-way ANOVA, where \*\*\**p* < 0.001, \*\**p* < 0.01.

**Supplementary Figure 3:**
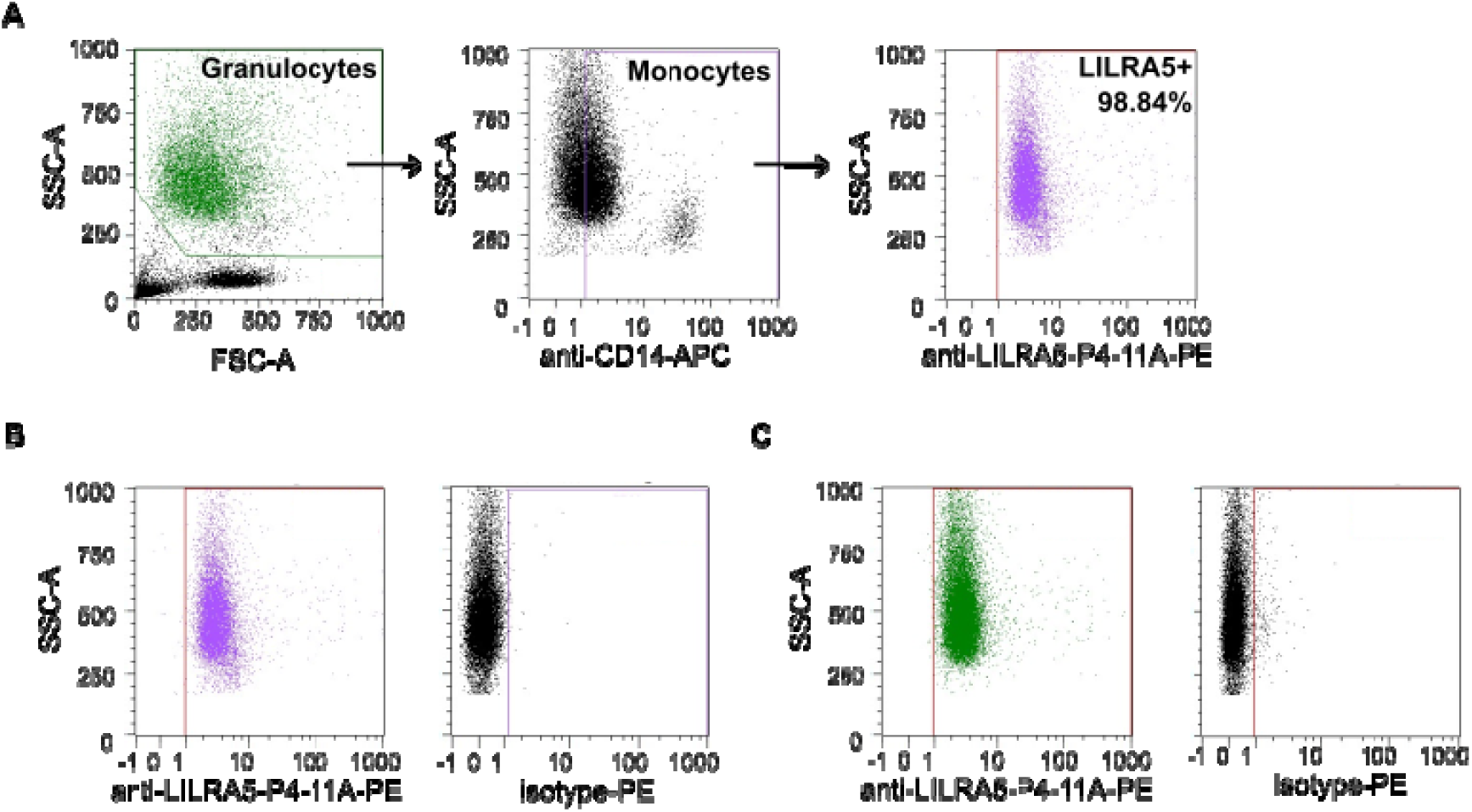
Gating strategy and expression of LILRA5 on monocytes. **(A)** Representative example showing the gating strategy used to identify monocytes in human whole blood using anti-CD14 and anti-LILRA5 (clone P4-11A). A first gate was set on physical parameters of SSC-A vs. FSC-A, then on SSC-A and SSC-H to eliminate doublets (not shown, but similarly presented in Fig 2), then monocytes and granulocytes were gated on CD14+ and CEACAM8+ (not shown), then on CD14+ events to identify monocytes. **(B)** Representative flow cytometry histogram showing LILRA5 expression on CD14+ monocytes. **(C)** Representative flow cytometry histogram showing LILRA5 expression on CEACAM8+ neutrophils.

**Supp. Fig 4.**
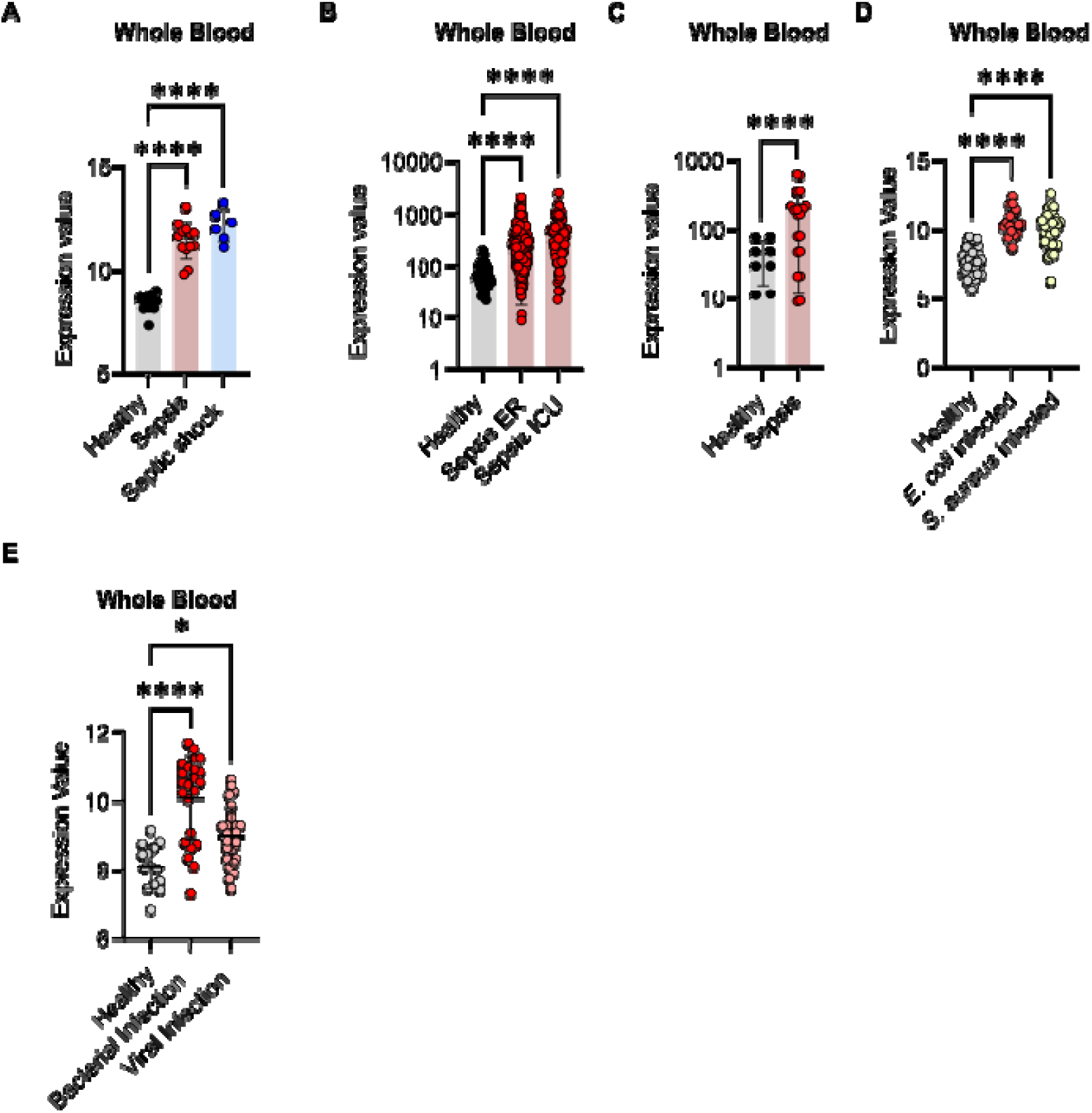
*LILRA5* expression during sepsis and systemic infection. **(A)** *LILRA5* expression in whole blood of healthy donors (*n* = 12) or sepsis patients (*n* = 13) or septic shock patients (*n* = 6) (GSE137342). Mean ± SD are shown. Statistics tested by *limma*, where adjusted \*\*\**p* < 0.001. **(B)** *LILRA5* expression in whole blood of healthy donors (*n* = 44) or sepsis patients presenting at emergency rooms (ER) (*n* = 266) or sepsis patients on intensive care units (ICU) (*n* = 82) (GSE185263). Mean ± SD are shown. Statistics tested by *limma*, where adjusted \*\*\**p* < 0.001. **(C)** *LILRA5* expression in whole blood of healthy donors (*n* = 8) or sepsis patients (*n* = 20) (GSE232753). Mean ± SD are shown. Statistics tested by *limma*, where adjusted \*\*\**p* < 0.001. **(D)** *LILRA5* expression in whole blood from healthy donors (*n* = 14), patients with bacterial infections (*n* = 24) and patients with viral infections (*n* = 28)(GSE72810). Mean ± SD are shown. Statistics tested by *limma*, where adjusted \*\*\**p* < 0.001. **(E)** *LILRA5* expression in whole blood from healthy donors (*n* = 43), patients with *E. coli* infection (*n* = 32), or patients with *S. aureus* infection (*n* = 19) (GSE33341). Mean ± SD are shown. Statistics tested by *limma*, where adjusted \*\*\**p* < 0.001.

## Acknowledgements

We thank Birgit Maranca-Hüwel (University of Duisburg-Essen), Carla J.C. de Haas and Piet Aerts (UMC Utrecht) for excellent technical support. We thank Bernhard B. Singer who passed away in January 2023 for his involvement in the design and experimental part of this work. This work was financed by the Biotechnology and Biological Sciences Research Council (BBSRC, UK) grant BB/V006495/1 to A.J.M.; the Wellcome Trust grant 225315/Z/22/Z to A.J.M.; by the European Union Horizon 2020 research and innovation programme under Grant Agreement 700862; and the German research foundation grant 316213987/C06/B08 and 550453157 to N.L. and A.T.P; and the German Research Foundation Excellence Strategy 390713860 to M.B.

## Author contributions

Z.F., M.R., I.K.-G., M.L., V.S., Y.Z., A.K. performed experiments. Z.F., M.R., I.K.-G., A.J.M analysed data, visualised data, and wrote the original draft. N.L., A.T.P., S.B, S.B.F and G.W. provided scientific help and critically reviewed the manuscript. A.J.M. I.K.-G. and B.B.S. conceptualised and directed the study. A.J.M. supervised the analysis and interpretation of the data. All authors contributed to the article and approved the submitted version.

## References

1. Dale DC, Boxer L, Liles WC. The phagocytes: neutrophils and monocytes. Blood. 2008 Aug 15;112(4):935–45.

2. Underhill DM, Rossnagle E, Lowell CA, Simmons RM. Dectin-1 activates Syk tyrosine kinase in a dynamic subset of macrophages for reactive oxygen production. Blood. 2005 Oct 1;106(7):2543–50.

3. Ravetch JV, Bolland S. IgG Fc receptors. Annu Rev Immunol. 2001;19:275–90.

4. Huizinga TW, van Kemenade F, Koenderman L, Dolman KM, von dem Borṣne AE, Tetteroo PA, et al. The 40-kDa Fc gamma receptor (FcRII) on human neutrophils is essential for the IgG-induced respiratory burst and IgG-induced phagocytosis. J Immunol. 1989 Apr 1;142(7):2365–9.

5. Lee WB, Kang JS, Yan JJ, Lee MS, Jeon BY, Cho SN, et al. Neutrophils Promote Mycobacterial Trehalose Dimycolate-Induced Lung Inflammation via the Mincle Pathway. PLoS Pathog. 2012 Apr 5;8(4):e1002614.

6. Brandsma AM, Hogarth PM, Nimmerjahn F, Leusen JHW. Clarifying the Confusion between Cytokine and Fc Receptor “Common Gamma Chain.” Immunity. 2016 Aug 16;45(2):225–6.

7. Hirayasu K, Arase H. Functional and genetic diversity of leukocyte immunoglobulin-like receptor and implication for disease associations. J Hum Genet. 2015 Nov;60(11):703– 8.

8. Mitchell A, Rentero C, Endoh Y, Hsu K, Gaus K, Geczy C, et al. LILRA5 is expressed by synovial tissue macrophages in rheumatoid arthritis, selectively induces pro-inflammatory cytokines and IL-10 and is regulated by TNF-alpha, IL-10 and IFN-gamma. Eur J Immunol. 2008 Dec;38(12):3459–73.

9. Borges L, Kubin M, Kuhlman T. LIR9, an immunoglobulin-superfamily-activating receptor, is expressed as a transmembrane and as a secreted molecule. Blood [Internet]. 2003 Feb 15 [cited 2023 Jun 8];101(4). Available from: https://pubmed.ncbi.nlm.nih.gov/12393390/

10. Lewis Marffy AL, McCarthy AJ. Leukocyte immunoglobulin-like receptors (LILRs) on human neutrophils: Modulators of infection and immunity. Front Immunol. 2020 May 13;11:857.

11. Burshtyn DN, Morcos C. The expanding spectrum of ligands for leukocyte Ig-like receptors. J Immunol. 2016 Feb 1;196(3):947–55.

12. Shang Y, Liu X, Wei L, Liang S, Zou Z, Wu M, et al. Leukocyte Immunoglobulin-like Receptor A5 Deletion Aggravates the Pathogenesis of Pseudomonas aeruginosa Keratitis by Promoting Proinflammatory Cytokines. Cornea. 2023 May 1;42(5):607–14.

13. Ning J, Fan X, Sun K, Wang X, Li H, Jia K, et al. Single-cell Sequence Analysis Combined with Multiple Machine Learning to Identify Markers in Sepsis Patients: LILRA5. Inflammation. 2023 Aug;46(4):1236–54.

14. Aleyd E, Heineke MH, van Egmond M. The era of the immunoglobulin A Fc receptor FcαRI; its function and potential as target in disease. Immunol Rev. 2015 Nov;268(1):123–38.

15. Zhao Y, van Woudenbergh E, Zhu J, Heck AJR, van Kessel KPM, de Haas CJC, et al. The Orphan Immune Receptor LILRB3 Modulates Fc Receptor–Mediated Functions of Neutrophils. J Immunol. 2020 Feb 15;204(4):954–66.

16. Gao XM, Zhou XH, Jia MW, Wang XZ, Liu D. Identification of key genes in sepsis by WGCNA. Prev Med. 2023 Jul;172:107540.

17. Demaret J, Venet F, Friggeri A, Cazalis MA, Plassais J, Jallades L, et al. Marked alterations of neutrophil functions during sepsis-induced immunosuppression. J Leukoc Biol. 2015 Dec;98(6):1081–90.

18. Dix A, Hünniger K, Weber M, Guthke R, Kurzai O, Linde J. Biomarker-based classification of bacterial and fungal whole-blood infections in a genome-wide expression study. Front Microbiol. 2015 Mar 11;6:171.

19. Herberg JA, Kaforou M, Wright VJ, Shailes H, Eleftherohorinou H, Hoggart CJ, et al. Diagnostic Test Accuracy of a 2-Transcript Host RNA Signature for Discriminating Bacterial vs Viral Infection in Febrile Children. JAMA. 2016;316(8):835–45.

20. Herwanto V, Tang B, Wang Y, Shojaei M, Nalos M, Shetty A, et al. Blood transcriptome analysis of patients with uncomplicated bacterial infection and sepsis. BMC Res Notes. 2021 Feb 27;14(1):76.

21. Futosi K, Fodor S, Mócsai A. Neutrophil cell surface receptors and their intracellular signal transduction pathways. Int Immunopharmacol. 2013 Nov;17(3):638–50.

22. Jones DC, Roghanian A, Brown DP, Chang C, Allen RL, Trowsdale J, et al. Alternative mRNA splicing creates transcripts encoding soluble proteins from most LILR genes. Eur J Immunol. 2009 Nov;39(11):3195–206.

23. Giraudi PJ, Pascut D, Banfi C, Ghilardi S, Tiribelli C, Bondesan A, et al. Serum proteome signatures associated with liver steatosis in adolescents with obesity. J Endocrinol Invest [Internet]. 2024 Jul 17; Available from: 10.1007/s40618-024-02419-x

24. Fu YL, Harrison RE. Microbial Phagocytic Receptors and Their Potential Involvement in Cytokine Induction in Macrophages. Front Immunol. 2021 Apr 29;12:662063.

25. Hirayasu K, Saito F, Suenaga T, Shida K, Arase N, Oikawa K, et al. Microbially cleaved immunoglobulins are sensed by the innate immune receptor LILRA2. Nature microbiology [Internet]. 2016 Apr 25 [cited 2023 Nov 15];1(6). Available from: https://pubmed.ncbi.nlm.nih.gov/27572839/

26. van Sorge NM, Bonsor DA, Deng L, Lindahl E, Schmitt V, Lyndin M, et al. Bacterial protein domains with a novel Ig-like fold target human CEACAM receptors. EMBO J. 2021 Apr 1;40(7):e106103.

27. Köhler G, Milstein C. Continuous cultures of fused cells secreting antibody of predefined specificity. Nature. 1975 Aug 7;256(5517):495–7.

28. Ohtsuka M, Arase H, Takeuchi A, Yamasaki S, Shiina R, Suenaga T, et al. NFAM1, an immunoreceptor tyrosine-based activation motif-bearing molecule that regulates B cell development and signaling. Proc Natl Acad Sci U S A. 2004 May 25;101(21):8126–31.

29. Scicluna BP, Uhel F, van Vught LA, Wiewel MA, Hoogendijk AJ, Baessman I, et al. The leukocyte non-coding RNA landscape in critically ill patients with sepsis. Elife [Internet]. 2020 Dec 11;9. Available from: 10.7554/eLife.58597

30. Baghela A, Pena OM, Lee AH, Baquir B, Falsafi R, An A, et al. Predicting sepsis severity at first clinical presentation: The role of endotypes and mechanistic signatures. EBioMedicine. 2022 Jan;75(103776):103776.

31. Hortová-Kohoutková M, De Zuani M, Lázničková P, Bendíčková K, Mrkva O, Andrejčinová I, et al. Polymorphonuclear Cells Show Features of Dysfunctional Activation During Fatal Sepsis. Front Immunol. 2021 Dec 13;12:741484.

32. Ahn SH, Tsalik EL, Cyr DD, Zhang Y, van Velkinburgh JC, Langley RJ, et al. Gene expression-based classifiers identify Staphylococcus aureus infection in mice and humans. PLoS One. 2013 Jan 9;8(1):e48979.

33. Song R, Gao Y, Dozmorov I, Malladi V, Saha I, McDaniel MM, et al. IRF1 governs the differential interferon-stimulated gene responses in human monocytes and macrophages by regulating chromatin accessibility. Cell Rep. 2021 Mar 23;34(12):108891.

34. Kleinertz H, Hepner-Schefczyk M, Ehnert S, Claus M, Halbgebauer R, Boller L, et al. Circulating growth/differentiation factor 15 is associated with human CD56 natural killer cell dysfunction and nosocomial infection in severe systemic inflammation. EBioMedicine. 2019 May;43:380–91.

35. Singer M, Deutschman CS, Seymour CW, Shankar-Hari M, Annane D, Bauer M, et al. The Third International Consensus Definitions for Sepsis and Septic Shock (Sepsis-3). JAMA [Internet]. 2016 Feb 23 [cited 2023 Jun 12];315(8). Available from: https://pubmed.ncbi.nlm.nih.gov/26903338/

